# Combined pigment and metatranscriptomic analysis reveals synchronized diel patterns of phenotypic light response across domains in the open ocean

**DOI:** 10.1101/2020.05.12.091322

**Authors:** Kevin W. Becker, Matthew J. Harke, Daniel R. Mende, Daniel Muratore, Joshua S. Weitz, Edward F. DeLong, Sonya T. Dyhrman, Benjamin A.S. Van Mooy

## Abstract

Sunlight is the most important environmental control on diel fluctuations in phytoplankton activity, and understanding diel microbial processes is essential to the study of oceanic biogeochemical cycles. Yet, little is known about the *in situ* frequency of phytoplankton metabolic activities and their coordination across different populations. We investigated the diel orchestration of phytoplankton activity involved in photosynthesis, photoacclimation, and photoprotection by analyzing the pigment and quinone distribution in combination with metatranscriptomes in the surface waters of the North Pacific Subtropical Gyre (NPSG). We found diel cycles in pigment abundances resulting from the balance of their synthesis and consumption. The night represents a metabolic recovery phase to refill cellular pigment stores, while the photosystems are remodeled towards photoprotection during the day. Transcript levels of genes involved in photosynthesis and pigment metabolism had highly synchronized diel expression patterns among all taxa, suggesting that there are similar regulatory mechanisms for light and energy metabolism across domains, and that other environmental factors drive niche differentiation. Observed decoupling of diel oscillations in transcripts and related pigments in the NPSG indicates that pigment abundance is modulated by environmental factors extending beyond gene expression/regulation, showing that metatranscriptomes may provide only limited insights on real-time photophysiological metabolism.

## Introduction

Sunlight regulates the growth and cellular processes of all photosynthetic organisms in the ocean. It plays a pivotal role for ocean ecosystem processes, many of which are driven by energy provided by photosynthesis. The conversion of light into chemical energy is achieved by the absorption of photons by pigments in light-harvesting complexes followed by the transfer of that energy to the reaction center, where it is used to initiate the electron transfer and charge separation processes. Plastoquinones mediate the transfer of electrons between photosystem II (PSII), the cytochrome *b*_6_f complex (Cyt*b*_6_f), and photosystem I (PSI). The net result is the generation of ATP and NADPH, which are then used for carbon fixation, biosynthesis, and aerobic respiration (see review by Eberhard et al. [1]). Pigments play a central role in energy transport within light-harvesting complexes and photosystems of all photosynthetic organisms (Fig. 1). In particular, chlorophyll (Chl) is vital in these processes, performing both light harvesting and electron transfer, and it is typically the most abundant pigment. Carotenoid pigments also participate in energy transport and have an additional role in photoacclimation.

**Fig. 1.**
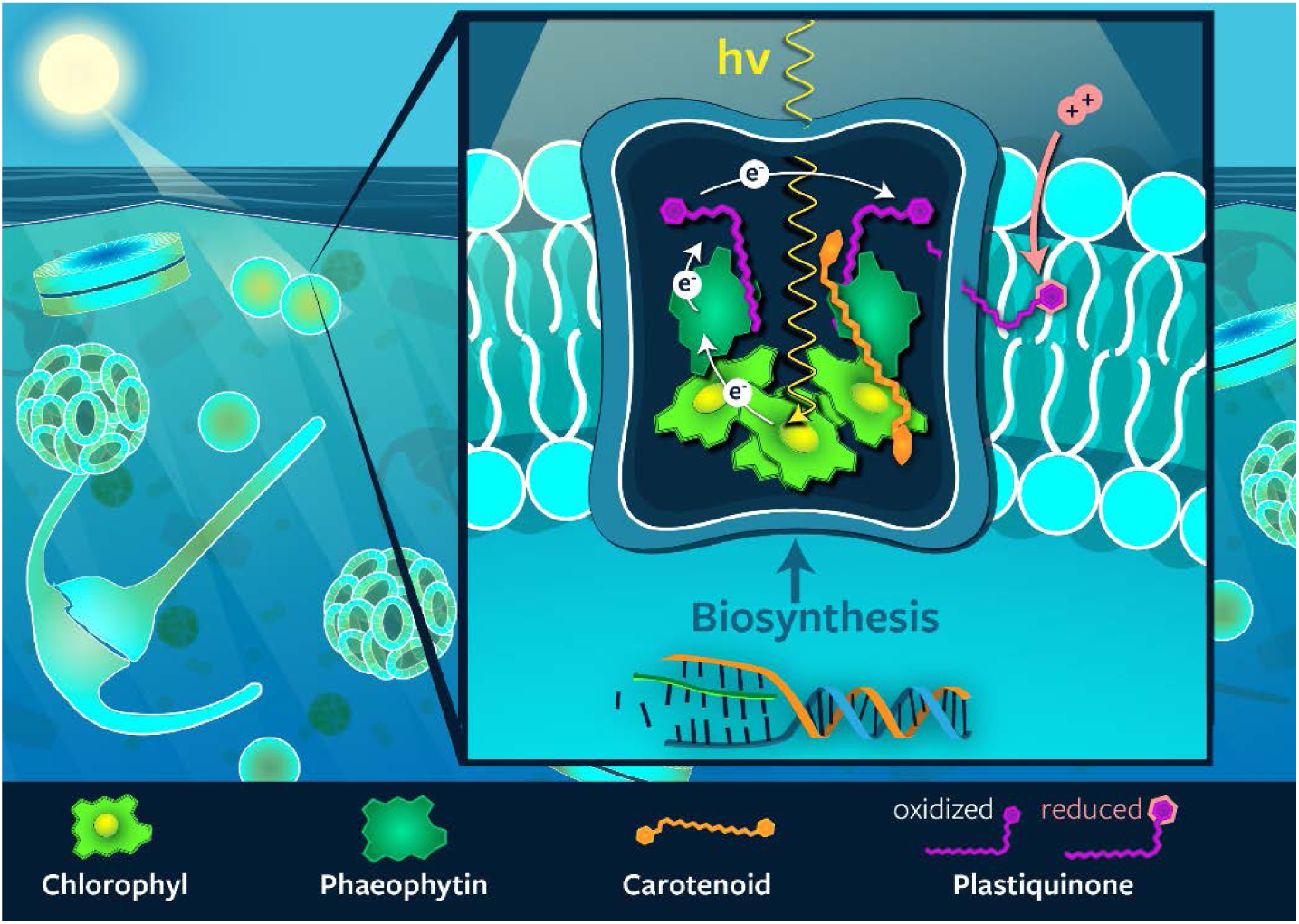
Schematic representation of the structure of photosystem II in photosynthetic membranes. The relative locations of the major pigments and quinones as well as the pathway of electrons through these molecules is shown.

Phytoplankton have developed a wide variety of pigment-based mechanisms for adapting to the variable light field experienced in the upper ocean. These mechanisms include downsizing of the photosynthetic antenna size by reducing the pool size of Chl, using xanthophyll cycling to arrest photoinhibition, and quenching singlet oxygen by carotenes [2]. Pigment molecules thus play key roles in the acclimation (short-term) and adaptation (long-term) of phytoplankton to the variability in light. In addition to their physiological importance, evolutionary divergence has led to the formation of a variety of chlorophyll and carotenoid pigments in different photosynthetic organisms, which allows us to delineate activity amongst phytoplankton groups [e.g., 3]. Also, the biosynthetic pathways of pigments are well known [e.g., 4]. This allows investigation of expression patterns of chlorophyll- and carotenoid pigment-related genes to gain further insight into the activity of individual phytoplankton groups, enabling taxonomic resolution not necessarily provided by pigment analysis alone.

Diel oscillations in metabolic activities have been observed for both gene expression [e.g., 5–8] and pigment profiles [e.g., 9, 10] in natural phytoplankton populations. However, there has been no study of the relationship between phytoplankton pigments and photosynthesis-related gene expression over the diel cycle. Little is known about the synchronicity of photosynthetic metabolic pathways between different phytoplankton groups in nutrient deplete marine environments such as the subtropical gyres. Yet, the nutrient scarcity requires niche and/or resource partitioning by different phytoplankton to facilitate growth in the face of competition [11]. Additionally, temporal anomalies of phytoplankton biomass and cellular pigment content impact the optical properties of the surface ocean influencing the interpretation of satellite-derived ocean color observations and global models using these data [12]. In these studies, diel oscillations are typically not considered because satellite observations take place only at a narrow interval of the diurnal cycle. Instead, most global ocean ecosystem models based on satellite data assume that the observed phytoplankton communities are under steady state growth conditions with respect to daily time-scales [e.g., 13]. Thus, better understanding group-specific differences as well as bulk community acclimation over the diel cycle is important for informing models relying on photoacclimation parameters to determine growth and accumulation rates as well as primary production [12, 14].

In this study, we used a combined analysis of pigments, quinones, and metatranscriptomes to investigate the diel orchestration of phytoplankton activity involved in photosynthesis, photoacclimation, and photoprotection in the surface waters of the North Pacific Subtropical Gyre (NPSG). An intensive multidisciplinary field campaign near station ALOHA in summer 2015 made use of a Lagrangian sampling strategy that mitigates the effect of spatial variability and allowed us to investigate microbial populations within the same water parcel over time [15]. In light of the importance of the subtropical gyres of the oceans for global biogeochemical cycles [16, 17], it is crucial to know how diel cycles of photoacclimation in phytoplankton affect the pigment composition in these ecosystems.

## Results and discussion

### Diel oscillations of chloropigments and carotenoids

The majority of detected pigment and quinone molecules investigated here displayed clear, statistically significant diel oscillations in abundance based on the Rhythmicity Analysis Incorporating Non-parametric Methods (RAIN) hypothesis-testing method ([15, 18, 19]; Supplementary Table 1). Diel oscillations of chloropigments, i.e., mono- and divinyl-Chls, showed maximum concentrations at dawn (6:00 h local time) and minimum concentrations during daylight hours (Fig. 2; Fig. S1). This pattern is distinct from previous investigations of diurnal Chl, which showed a mid-day maximum and a night-time minimum especially in data integrated through the water column [10, 20, 21]. Based on these previous observations, it was thought that Chl synthesis directly reflects primary production by photoadaptation, with increasing cellular chlorophyll during the day and chlorophyll break down (or dilution by cell division) at night [e.g., 22]. Alternatively, mid-day minima in surface water Chl that have frequently been observed in data from *in vivo* Chl fluorescence have been interpreted to result from non-photochemical quenching, not absolute decreases in Chl concentrations. The reasoning behind this interpretation is that variability in quantum yield for fluorescence is significant and complicates the interpretation of Chl estimates from in vivo fluorescence at high light intensities (e.g., [23–25]). However, here we used mass spectrometry to determine Chl concentrations from lipid extracts, which is insensitive to quenching. Other studies, measuring extracted Chl, also showed clear and reproducible night-time maxima for both near surface waters (e.g., [9]) and culture experiments at high light intensities (e.g., [26–29]). Additionally, Fouilland et al. [30] found a depth-dependence of diel variations in Rubisco activity and Chl concentration per cell further supporting a light regime-dependence of Chl profiles.

**Fig. 2.**
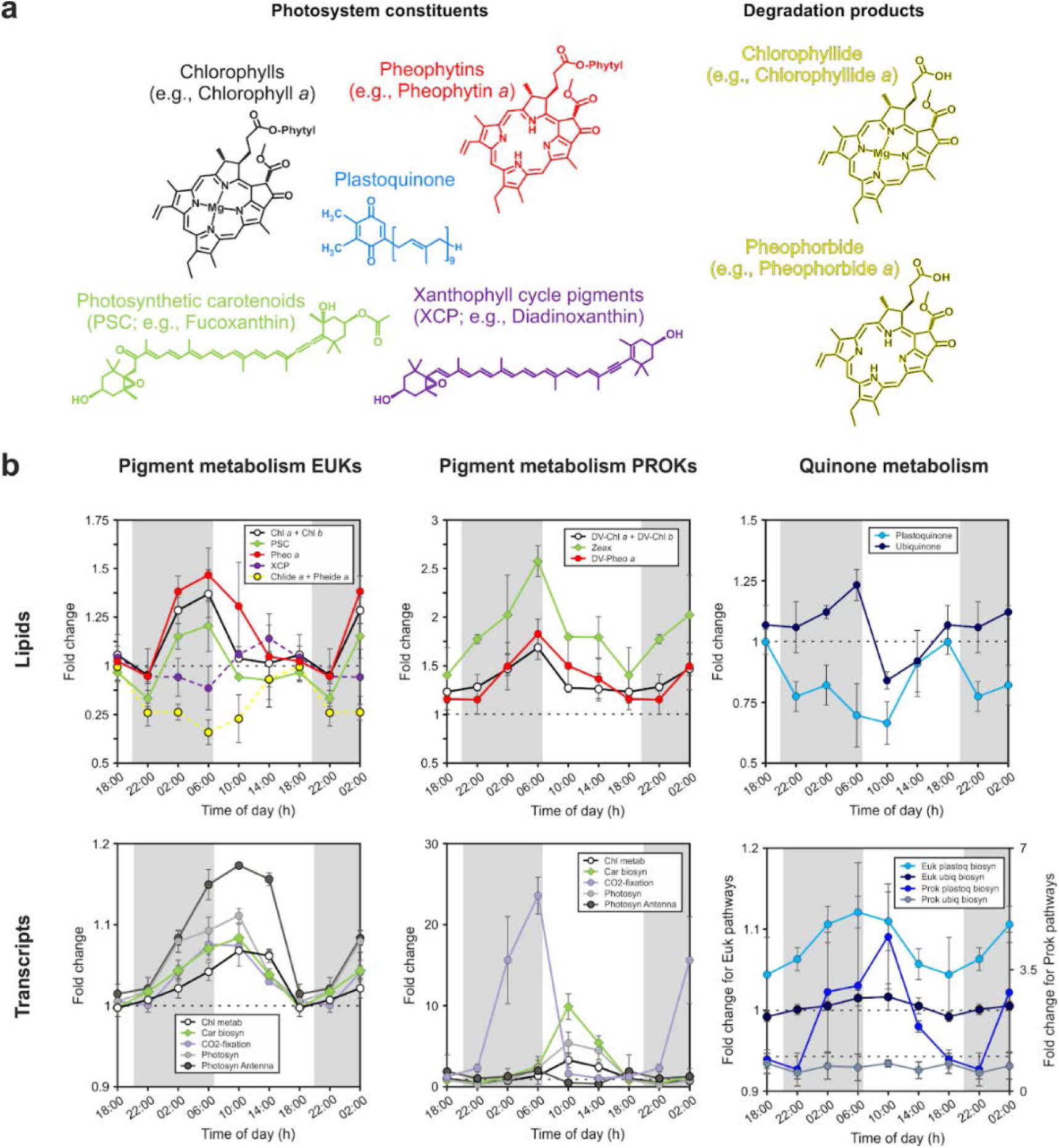
Diel oscillations of pigments and pigment-related transcripts. **a** Molecular structure of investigated pigments and respiratory quinones associated with the photosystem. **b** Time-of-day average of fold change relative to 18:00 h (local time) of chloro- and carotenoid pigments, quinones, as well as transcript associated with the metabolism of the pigment and quinone molecules. Error bars represent the standard deviation of averaged values for each time of day (n≥6). Molecular structures and pigment and quinone data are colored according to compound group. If colors for transcripts match those of pigments or quinones, transcripts are related to the metabolism of the respective molecule.

The photosynthetic carotenoids (PSC) aligned with Chl over the diel cycle (Fig. 2; Fig S1) and no periodic variations were observed in ratios among PSC pigments (Fig. S2), which has previously been observed for both phytoplankton cultures [31] and surface ocean samples [32] and is consistent with PSC and Chl having similar functions [33]. The most abundant PSC in surface waters were fucoxanthin (diatom biomarker), 19’-butanoyloxyfucoxanthin (pelagophyte biomarker), and 19’-hexanoyloxyfucoxanthin (prymnesiophyte biomarker). All showed similar trends, with a nocturnal increase leading to a pre-dawn maximum and a minimum later, during daylight hours (Fig. 2 and Fig. S2). Similar to Chls, the nocturnal increase may seem counterintuitive as light harvesting molecules are needed during the day for photosynthesis. However, at high photon flux, the rate of photon absorption by Chl antenna far exceeds the rate at which photons can be utilized for photosynthesis; to avoid over-excitation of the photosystem, high light acclimation typically involves downsizing of the antenna of both photosystems [34, 35], in particular of PSII [36]. The observed profiles thus likely represent the balance of photoprotective and photorepair mechanisms.

While the observed depression in Chl and PSC during daylight might be explained by PSII damage due to high-light stress, the increase at night clearly indicates synthesis in the dark. Increased vertical mixing during the night could also explain this signal, but the surface mixed layer typically ranges from ∼0-40 m [37], which is too shallow to transport large amounts of Chl, for example, from the deep Chl maximum (at ∼100 m at station ALOHA our sampling site and times [15, see Fig. 1 therein]) to the sampling depth of 15 m. Additionally, *Prochlorococcus* cell numbers remained constant over the diel cycle in our samples, which results in increasing divinyl-Chl/cell quotas during the night (Fig. S2) and supports the hypothesis of night-time synthesis. The energy and carbon for the *de novo* synthesis of Chls in the dark is likely provided by carbon stores, for example glycogen in cyanobacteria or triacylglycerols (TAGs) in eukaryotic phytoplankton. Consistent with this, we recently showed that nanoeukaryotic phytoplankton in the NPSG accumulate large amounts of TAGs during the day and subsequently consume them at night [38]. A similar mechanism has been proposed for dark synthesis of proteins in several phytoplankton (e.g., [39–41]) and our findings extend the potential utilization of C stores by phytoplankton for dark synthesis of pigments.

Pheophytin and divinyl-pheophytin profiles were tightly coupled to those of Chl and divinyl-Chl, respectively, with peak times at night (Fig. 2). While the presence of pheophytin in marine samples has traditionally been interpreted to reflect dead or dying cells due to the high abundance of these pigments in zooplankton guts and sinking particles, which are unlikely to be consistently captured in our small volume samples [42–46], we hypothesize that the pheophytins may in fact be associated with living phytoplankton. Most notably, pheophytin has been found to play an important role in electron transport in PSII by being the primary electron acceptor for excited Chl [47] and has been frequently detected in isolated PSII with a 6:2 stoichiometry of Chl and pheophytin [e.g., 48]. Additionally, pheophytin and Chl share a biochemical pathway, where Chl is converted to pheophytin by a magnesium dechelatase [49] supporting the idea of simultaneous dark synthesis of Chls and pheophytins. Other Chl related structures, pheophorbide and chlorophyllide, were out of phase with Chl and pheophytin, both showing peak times at 14:00 h (Fig. 2). These Chl related structures are not known to participate in photosynthetic reactions, suggesting daytime peaks are likely associated with photodamage and photoprotective mechanisms.

High light stress during daylight is also apparent from xanthophyll cycle pigments, essential molecules that channel excess energy away from chlorophylls for protection against photooxidative damage [50]. Diadinoxanthin (Dd) and diatoxanthin (Dt), which are part of the Dd/Dt xanthophyll cycle predominantly possessed by haptophytes, diatoms and dinoflagellates in the NPSG [51], were the most abundant xanthophyll cycle pigments detected (Fig. S1). The sum of Dt and Dd showed clear diel cycles with maximum abundances during the day (peak time at 14:00 h; Fig. 2). This pattern indicates that the cellular xanthophyll pigment pool increased during daylight hours likely due to photoadaptation triggered by a change of light intensity over the course of the day [32]. The de-epoxidation state (DES), which is defined as [Dt]/[Dt+Dd], showed a minimum during the day (Fig. S2), which contrasts observations from culture experiments [52] and the field [32]. This pattern is likely related to severe high-light stress at 15 m water depth where our samples were collected. Oxidative stress due to an increased formation of various ROS species consumes Dt by degrading it to low molecular weight compounds [53, 54], requiring re-synthesis for the pool to be refilled. Potentially, phytoplankton refill cellular Dt stores during the night. We therefore looked for Violaxanthin de-epoxidases, which convert Dd to Dt in the xanthophyll cycle [55], and find that transcript abundances for the diatoms and haptophytes were out of phase with XCPs (Fig. S10). Additionally, Zeaxanthin epoxidases, which participate in non-photochemical quenching by regulating the level of epoxy-free xanthophylls in photosynthetic energy conversion, showed statistically significant diel oscillation for diatoms and haptophytes with peak times during the day at 10:00 h local time (Fig. S11). Our results from the pigment analysis thus show that high light conditions in the upper photic zone results in severe photooxidative stress for phytoplankton, which is displayed in a distinct and different diel xanthophyll cycle pigment and Chl degradation product patterns compared to observations from the deeper photic zone [32]. Through this, phytoplankton are likely able to maintain the balance between dissipation and utilization of light energy to minimize the generation of ROS and resulting molecular damage. Since net primary production is typically invariable throughout the mixed layer at station ALOHA [56], the observed pigment signal represents a light adaptation signal as opposed to an increased photodegradation state of near surface phytoplankton cells.

### Diel patterns of isoprenoid quinones

Analyses of quinones in the marine water column are still relatively rare [38, 57, 58], but due to their physiological importance in electron transport in both photosynthesis and cellular respiration, quinones provide valuable information on microbial activities. While plastoquinones (PQs) are primarily involved in electron transport of PSII, ubiquinones (UQs) are associated with aerobic respiration [59]. Recent studies suggested PQs and UQs might additionally be involved in photoprotective mechanisms by acting as ROS scavengers [60–63]. In our samples, both quinone groups showed distinct and statistically significant diel patterns (Fig. 2; Table S1) with UQs peaking at the end of the night (6:00 h; cf. [38]) and PQs peaking at the end of the day (18:00 h, Fig. 2). These two groups of structurally similar molecules were thus out of phase by 12 h. The pre-dawn maximum of UQs is likely associated with enhanced respiration of storage lipids, which have been shown to accumulate during the day and are consumed at night [38]. The daytime increase of PQs may partly be driven by growth, but similar to other photosystem constituents, photoacclimation mechanisms play a major role for intercellular PQ abundances. Increasing the rates of electron transport processes can protect PSII when absorption of quanta exceeds the requirements of photosynthetic carbon metabolism [64], which, in turn, requires an increase in the PQ pool size and may in part be responsible for observed daytime maxima of PQs. Additionally, in plants, PQs have been shown to act as ROS scavengers under high-light stress [65, 66]. These processes deplete PQs due to photooxidation during daylight hours. Hence, they have to be re-synthesized to keep this important function. The daytime increase in the proportion of the PQ to chlorophyll pool (Fig. S2) is likely due to the different organization of chloroplast structure. In plants, chloroplasts have less antenna chlorophyll per electron transport chain under high light than low light [67].

### Diel oscillations of transcripts involved in photosynthesis

In addition to measuring diel oscillations in pigments and quinones, we further explored diel transcriptional rhythms of genes involved in photosynthesis and pigment metabolism of the dominant bacterial and eukaryotic groups at station ALOHA. Transcripts of all photosynthetic populations showed statistically significant diel oscillations for most investigated pathways (Supplementary Table 2). Almost all significant diel transcripts associated with photosynthesis and pigment metabolism peaked in the first part of the day (10:00 h; Fig. 3a), suggesting their transcriptional regulation in phototrophs is strongly synchronized across diverse taxa (within the temporal resolution of our measurements). These observations further imply that the pigment metabolism network is conserved across domains in the oligotrophic open ocean (see Figs. S3 – S9 for profiles of individual phytoplankton groups). This complements recent findings by Kolody et al. in coastal systems [8] who observed periodic photosynthesis-related transcripts that were conserved across diverse phototrophic lineages in a high-nutrient coastal environment. Thus, our data is consistent with a multispecies synchronous diel cycles of genes associated with photosynthesis and pigment metabolism with increasing transcript abundances during dark hours and peak times in the mid-morning. Nightly expression of photosynthesis genes in natural photoheterotrophic bacteria, followed by a drawdown shortly after light onset, has been suggested to be related to preparing cells for efficient solar energy harvest in the early morning hours [6]. Our data from cyanobacteria and eukaryotic phytoplankton and consistent with this hypothesis. A notable exception of this pattern was *Crocosphaera*. Transcripts for most pathways in this diazotroph only showed a weak relationship or no significant positive correlation with the pathways in the other phytoplankton populations. However, this is in agreement with other culture [68] and field-based observations [15, 69]. The process of N_2_ fixation in this unicellular cyanobacteria seems to require a tightly orchestrated diel cycle distinct from other diazotrophs and non-diazotrophic microorganisms [15, 69].

**Fig. 3.**
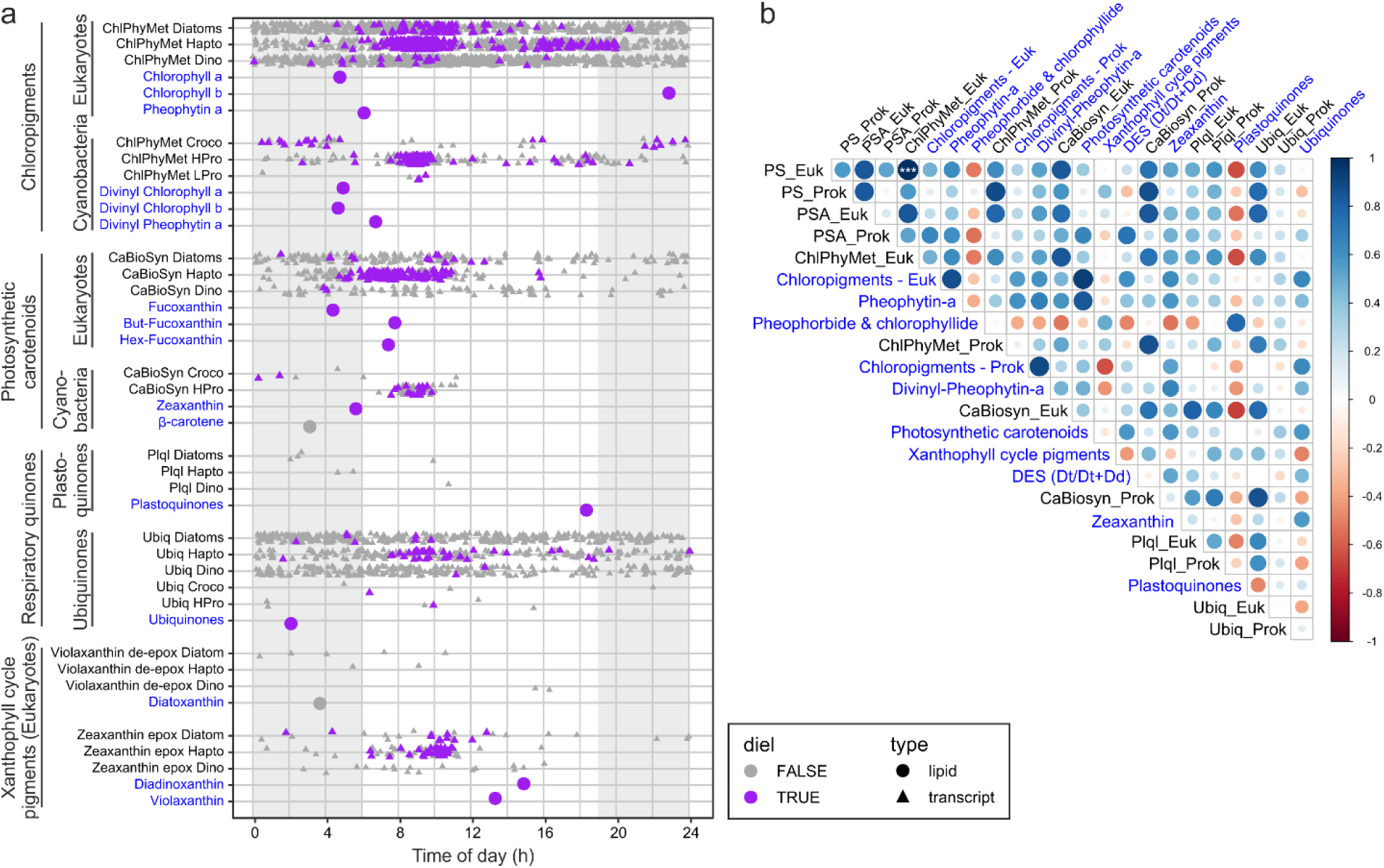
Relationship between periodic pigment-associated gene expression in phytoplankton transcripts and actual pigment abundances **a.** Peak times of expression of all transcripts assigned to KEGG pathways associated with pigment metabolism (triangles) and pigment molecule abundances (circles). Purple symbols denote transcripts or pigment molecules identified as significantly periodic (24-hour period) whereas gray symbols denote peak times without a significant diel component in peak expression or pigment abundance, respectively. **b.** Spearman’s rank correlation matrix of the different pigment and quinone classes (blue text) and aggregated genes within the KEGG pathways Photosynthesis (PS), Carotenoid biosynthesis (CaBiosyn), Chlorophyll and porphyrin metabolism (ChlPhyMet), Photosynthesis - antenna proteins (PSA), Plastoquinol biosynthesis (Plql), Ubiquinone and other terpenoid-quinone biosynthesis (Ubiq) separated into prokaryotes (Prok) and eukaryotes (Euk).

Strong diel rhythmicity in transcripts of eukaryotic phytoplankton including haptophytes and diatoms is expected from earlier culture and field experiments [e.g., 5, 8, 19, 70–72]. However, peak times for pathways involved in photosynthesis and pigment metabolism seem to be variable in culture experiments. For example, light harvesting antenna protein gene expression was enriched in late afternoon (2 to 6 pm) for iron-replete cultures of the diatom *Phaeodactylum tricornutum*, along with genes involved in porphyrin and chlorophyll metabolism [71], while enrichment for the KEGG pathways photosynthesis and porphyrin and chlorophyll metabolism was observed in the dawn for the diatom *Thalassiosira pseudonana* [70]. This variability in the timing of gene expression contrasts our findings from natural phytoplankton populations in the NPSG, which showed synchronized expression patterns across diverse phytoplankton. This synchrony across taxa has recently also been shown for a coastal environment [8] and together the results indicate that responses to signals in the environment, possibly including organismal interactions, play an important role in shaping gene expression patterns of natural phytoplankton populations.

### Decoupling between transcription and pigment production

The combination of pigment analysis and metatranscriptomics further shed light on the complex interplay between metabolic pathways and actual metabolite concentrations. In a previous study of energy storage mechanisms of natural phytoplankton populations at station ALOHA, we showed that gene expression and lipid abundances were decoupled - despite distinct diel patterns in storage lipid abundance, transcript abundance for the genes involved in the final step of their biosynthesis remained relatively constant over the diel cycle [38; Fig 4]. In addition, the correlations we observe between transcripts and lipids (shown in Fig. 3b) are not significantly stronger than the correlations expected from comparing independent time series (see Fig. S13). In culture experiments with the marine diatom *Phaeodactylum tricornutum*, during high light acclimation, Chl metabolism at the transcriptional level was down-regulated at an early stage while Chl *a* concentrations showed a lag time to this initial transcriptional response [73], which is what would be expected for an ideal system. However, the change of concentrations of pigments or any metabolite is governed by the difference between production and consumption. For example, the observed offset in peak time between transcripts (mid-morning) and associated pigments (dawn, Fig. 3a) suggests the influence of additional regulatory processes for chlorophyll and carotenoid pigment abundances. Irradiance at 15 m at station ALOHA (700 µmol m^-2^ s^-1^) was higher than in the “high light” experiments conducted by Nymark et al. [73] (500 µmol m^-2^ s^-1^) and the different patterns between the culture experiments and our field based observations may be related to photoacclimation mechanisms. Our observations that transcripts peak approximately 4 hours later than their corresponding pigments might indicate that phytoplankton actively synthesize Chls and PSC at a high rate, but catabolic processes, i.e., a shift towards photoprotection, consume more molecules than are produced. Our data thus show that metatranscriptome data should not be interpreted as a real-time readout of photophysiological metabolism.

**Fig. 4.**
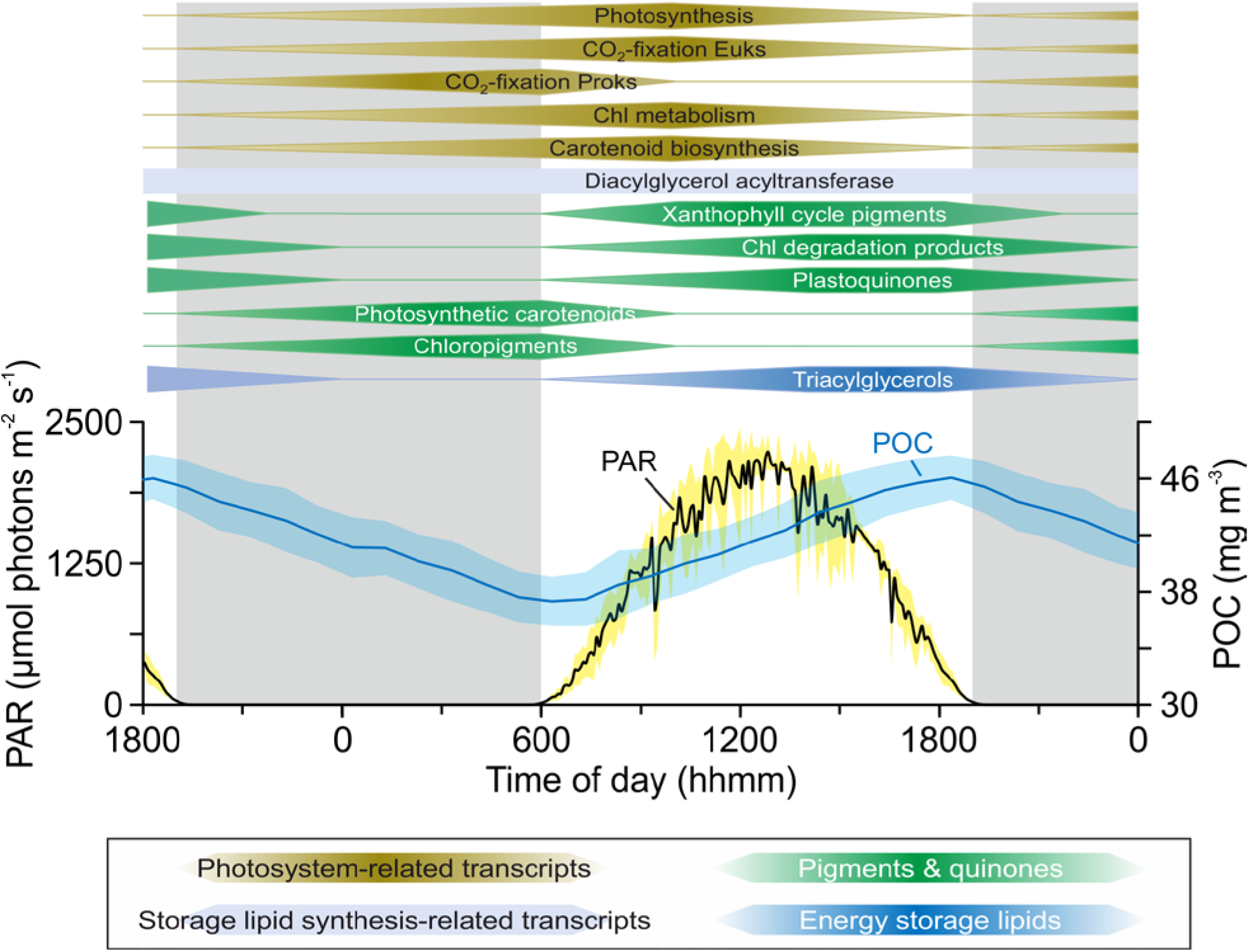
Conceptualized temporal separation of phytoplankton pigment, plastoquinone and transcript abundances during the dark-light cycle in surface waters (15 m) at station ALOHA. The timing was set based on Fig. 2 with the center of the polygon indicating the approximate peak time. Additionally, the particulate organic carbon (POC) and Photosynthetically Available Radiation (PAR) profiles averaged for one day/night cycle are shown. Surface PAR data was obtained from the HOT program database (http://hahana.soest.hawaii.edu/hoelegacy/hoelegacy.html), POC data from White et al. [76] and TAG (Triacylglycerols & Diacylglycerol acyltransferase) data from Becker et al. [38].

The response of cells towards ambient light conditions involves optimization of their photosystems in such a way that energy generation and utilization are in balance. Kana et al. [74] proposed that pigment abundance is modulated by environmental factors extending beyond gene expression/regulation, i.e., the initial response to high light intensities is self-adjusting. The light sensor that in a first step regulates photoacclimation in phytoplankton has been suggested to be the redox state of the quinone pool [74]. Although our methods do not allow direct determination of the redox state of the PQ pool because plastoquinols become spontaneously oxidized to plastoquinones when exposed to oxygen during sample analysis, we found significant diel oscillations in the expression of Chl *a*/*b* binding protein (*cab*) genes (Fig. S12). The *cab* genes are important for both light-harvesting and photoprotection in eukaryotic phytoplankton and are under transcriptional control by the redox poise of the PQ pool [75]. In cultures, under high light, it has been shown that the PQ pool becomes more reduced resulting in a suppression of *cab* mRNA synthesis and light harvesting complex production, and an ultimate decrease in cellular photosynthetic pigments [75]. We observed a decline in *cab* transcript abundances during daylight hours (Fig. S12) suggesting that this light intensity-dependent photon-sensing system is active in the NPSG. This process then likely triggers the down-regulation of photosynthesis and pigment metabolism related genes to keep pigments levels consistently low to prevent over-excitation of the photosystems at the high irradiance levels phytoplankton experience in the surface waters of the NPSG. Besides the insights our data provided on the (de)coupling between transcriptional and metabolite rhythms, our data suggest that the night reflects a metabolic recovery phase that is used by all photosynthetic organisms across domains to be prepared for photosynthetic reaction as soon as the sun rises, make optimal use of sunlight energy and grow efficiently.

## Conclusions

We are just beginning to understand in detail the mechanisms by which marine phytoplankton are able to maintain homeostasis and coordinate growth under metabolic and energetic shifts driven by the perpetual rising and setting sun. Our work demonstrates that combined lipidomics (here pigment and quinone profiles) and transcriptomics can provide mechanistic insights into rapid (i.e., sub-daily) microbial processes (Fig. 4). While pigments involved in light harvesting and energy transfer peaked at dawn, photoprotective pigments and Chl degradation products peaked later in the day. This succession of pigment metabolism according to their function suggests a recovery of cellular Chl and PSC stores at night and a dominance of photoprotective mechanisms during daylight. The diel cycles of combined pigment, quinone and transcriptomics data from the NPSG thus highlight the fundamental connection between sunlight and phytoplankton photosynthetic metabolism [74]. Furthermore, diel metabolic cycles were similar across all major phytoplankton taxa with few exceptions. Thus, harvesting light energy is not the basis for temporal niche differentiation among these taxa, which stands to reason because photons are in excess at 15 m depth and therefore are not a resource that phytoplankton compete with one another to obtain. Niche differentiation among phytoplankton occurs for processes involving competitive and growth-limiting substrates, such as nutrients, which are scarce in the surface of the oligotrophic ocean. Yet the daily synthesis of pigments – and the machinery required for this synthesis – place considerable burden on phytoplankton for growth-limiting substrates. Thus, any temporal partitioning in accessing growth-limiting substrates must also be accompanied by different modes for their storage, which are rarely considered [77, 78]. In summary, the data presented herein advances our knowledge of this fundamental process within photosynthetic microbes in the mixed layer, providing valuable detailed observations of transcriptional and pigment dynamics across kingdoms. These data will thus prove useful for future models linking remotely sensed ocean color to temporal dynamics of pigment concentrations or photosynthetic activity.

## Materials and methods

### Sample collection and lipid analysis

Seawater samples were collected during R/V Kilo Moana cruise KM1513 (July/August 2015) near Station ALOHA (22°45’N, 158°00’W) in the oligotrophic North Pacific Subtropical Gyre using standard Niskin-type bottles attached to a CTD rosette. Samples were collected every 4 h from 2 p.m. (local time) July 27, 2015 to 6 a.m. (local time) on July 30, 2015. Samples (∼2 L) were filtered using vacuum filtration (ca. −200 mm Hg) onto 47 mm diameter 0.2 µm hydrophilic Durapore filters (Millipore). Samples were immediately flash-frozen and stored at −196 °C until processing.

Lipids were extracted using a modified Bligh and Dyer protocol [79] with DNP-PE-C_16:0_/C_16:0_-DAG (2,4-dinitrophenyl phosphatidylethanolamine diacylglycerol; Avanti Polar Lipids, Inc., Alabaster, AL) used as an internal standard. Filter blanks were extracted and analyzed alongside environmental samples. The total lipid extract was analyzed by reverse phase high performance liquid chromatography (HPLC) mass spectrometry (MS) on an Agilent 1200 HPLC coupled to a Thermo Fisher Exactive Plus Orbitrap high resolution mass spectrometer (ThermoFisher Scientific, Waltham, MA, USA). HPLC and MS conditions are described in detail in [38] and [80]; modified after [81].

For identification and quantification of pigments and quinones, we used LOBSTAHS, an open-source lipidomics software pipeline based on adduct ion abundances and several other orthogonal criteria [80]. Pigments and quinones identified using LOBSTAHS were quantified from MS data after pre-processing with XCMS [82] and CAMERA [83]. XCMS peak detection was validated by manual identification using retention time as well as accurate molecular mass and isotope pattern matching of proposed sum formulas in full-scan mode and tandem MS (MS^2^) fragment spectra of representative compounds [38]. For validation of accuracy and reliability of LOBSTAHS identification and quantification, quality control (QC) samples of known composition and spiked lipid standards were interspersed with samples as described previously [80]. Lipid abundances obtained from LOBSTAHS were corrected for relative response of commercially available standards. Abundances of quinones were corrected for the response of a ubiquinone (UQ_10:10_) standard, chlorophylls and their associated compounds using a chlorophyll *a* standard, and carotenoid pigments using a *β*-carotene standard. All standards were purchased from Sigma Aldrich (St. Louis, MO, USA). Individual response factors were obtained from external standard curves by triplicate injection of a series of standard mixtures in amounts ranging from 0.15 to 40 pmol on column per standard. Data were corrected for differences in extraction efficiency using the recovery of the DNP-PE internal standard (Avanti Polar Lipids, Inc., Alabaster, AL; USA).

### Eukaryotic metatranscriptome analysis

Samples for the >5 μm microeukaryote metatranscriptomes were collected at the same time as lipid samples following Harke et al. [72]. Briefly, 20 L of seawater was collected in acid-washed carboys every 4 h from 10 p.m. (local time) July 26, 2015 to 6 a.m. (local time) on July 30, 2015 for a total of 21 time points. Seawater was prescreened through a 200 μm nylon mesh and then filtered onto two 5 μm polycarbonate filters (47 mm) via peristaltic pump, passing ∼10 L across each filter. Samples were then flash frozen in liquid N until extraction. Total RNA was extracted from individual filter sets (n = 2 per timepoint) using Qiagen RNeasy Mini Kit (Qiagen, Hilden, Germany) as in Harke et al. [72] and then sequenced on an Illumina HiSeq 2000 at the JP Sulzberger Genome Center (CUGC) using center protocols. PolyA-selected samples were sequenced to a depth of 90 million 100 bp, paired-end reads. Raw sequence quality was visualized with FastQC (https://www.bioinformatics.babraham.ac.uk/projects/fastqc/) and cleaned and trimmed using Trimmomatic [84] version 0.27 (paired-end mode, 4 bp-wide sliding window for quality below 15, minimum length of 25 bp). Processed reads were mapped to a reference database after Alexander et al. [85], constructed from the Marine Microbial Eukaryotic Transcriptome Sequencing Project (MMETSP; [86]. Transcripts within the reference database were annotated with KEGG using UProC [87]. Mapping was conducted with the Burrows-Wheeler Aligner (BWA-MEM, parameters –k 10 –aM; [88]) and read counts generated with the HTSeq 0.6.1 package (options –a 0, --m intersection-strict, -s no; [89]). Read counts were filtered for contigs with average read counts ≥ 10 across the time series and then normalized with DESeq2 “varianceStabilizingTransformation” command [90]. These environmental sequence data are deposited in the Sequence Read Archive through the National Center for Biotechnology Information under accession no. SRP136571, BioProject no. PRJNA437978. To facilitate comparisons with pigment and quinone data, transcripts occurring in dinoflagellate, haptophyte, and diatom taxa were mined for the following KEGG pathways: Carotenoid biosynthesis [PATH:ko00906], Porphyrin and chlorophyll metabolism [PATH:ko00860], Photosynthesis [PATH:ko00195], Carbon fixation in photosynthetic organisms [PATH:ko00710], and Ubiquinone and other terpenoid-quinone biosynthesis [PATH:ko00130]. In addition, signals involved in plastoquinol biosynthesis were separated out from Ubiquinone and other terpenoid-quinone biosynthesis to mirror pigment and quinone data.

### Prokaryotic metatranscriptome analysis

For this study, a previously published dataset of transcriptomes was used [15, 19], which consisted of bacterioplankton samples collected every four hours over the same study period. Sampling was performed as follows: 2 L of seawater were filtered onto 25 mm, 0.2 μm Supor PES Membrane Disc filters (Pall) using a peristaltic pump. The filtration time was between 15 and 20 min and filters were placed in RNALater (Ambion) immediately afterwards, and preserved at −80 °C until processing. Molecular standard mixtures for quantitative transcriptomics were prepared as previously described [91], and, 50 μl of each standard group was added to sample lysate before RNA purification. Metatranscriptomic libraries were prepared for sequencing with the addition of 5–50 ng of RNA to the ScriptSeq cDNA V2 library preparation kit (Epicentre). Metatranscriptomic samples were sequenced with an Illumina NextSeq 500 system using V2 high output 300 cycle reagent kit with PHIX control added. Reads were mapped to the station ALOHA gfne catalog [92] using LAST [93]. Transcripts were quantified through normalization of raw hit counts using molecular standards [15].Transcripts in *Prochlorococcus* and *Crocosphaera* were mined for the same KEGG pathways as the eukaryotes.

### Statistical analysis

The statistical significance of diel oscillations of pigment, quinone and transcript abundances was tested using the RAIN package in R [see 15, 18, 19]. Resulting p-values from RAIN analysis were corrected for false discovery using the ‘padjust’ function in R with Benjamini-Hochberg method [94]. Corrected *p*-values ≤0.05 were considered to have significant diel periodicity. Peak times were calculated with a harmonic regression, fitting the expression data to a sine curve.

To evaluate rank correlations between pigment and transcript time series, data were centered to a mean of zero and scaled to one standard deviation to facilitate inter-comparability between data with different units. Then, pairwise spearman rank correlations (spearman’s rho) were calculated between all pairs of measured time series. To address autocorrelation, we evaluated whether observed correlations were stronger than independent identically distributed random walks, which are known to often be spuriously correlated [95]. To test this, the differences between consecutive observations were calculated for all measured time series. The frequency distributions of observed differences were then used to bootstrap sample 20 random differences and cumulatively summed to simulate a random walk. This process was repeated and we bootstrap sampled 100,000 such random walks independently to create a null ensemble of time series with similar autocorrelation-1 structure to the data. Bootstrap sampling was then used to randomly select 100,000 pairs of random walks and rank correlations were calculated to generate a Monte Carlo simulation of the distribution of spearman’s rho between independent identically distributed random walks with step sizes similar to those observed in the data (Fig. S13). This distribution was then used to calculate empirical one-tailed p-values of observed rank correlations. The adaptive Benjamini-Hochberg method was applied to account for multiple testing [94]. All calculations were carried out using the R statistical computing language v 3.6.3 with functionalities from the ‘tidyverse’ set of packages version 1.2.1.

## Acknowledgement

We are grateful to the officers and crew of the R/V Kilo Moana cruise KM1513 (HOE Legacy II). We thank Helen F. Fredricks for help with lipidomics analysis, and Daniel J. Repeta for help with onboard sample collection. This work was funded by a grant from the Simons Foundation (SCOPE, Award # 329108, B.A.S.V.M. S.T.D, E.F.D.), and Gordon and Betty Moore Foundation (grant #3777, E.F.D.). K.W.B. was further supported by the Postdoctoral Scholarship Program at Woods Hole Oceanographic Institution & U.S. Geological Survey.

## Author contributions

B.A.S.V.M., S.T.D., E.F.D. conceived the study. K.W.B., M.J.H., and D.R.M. conducted and analyzed lipidomic, eukaryotic metatranscriptomic and prokaryote transcriptomic studies, respectively. J.S.W. and D.M. supported data analysis. K.W.B. and M.J.H. wrote the paper with contributions from all authors.

