## Supplemental Figures S1-S13 for "Combined pigment and metatranscriptomic analysis reveals synchronized diel patterns of phenotypic light response across domains in the open ocean"

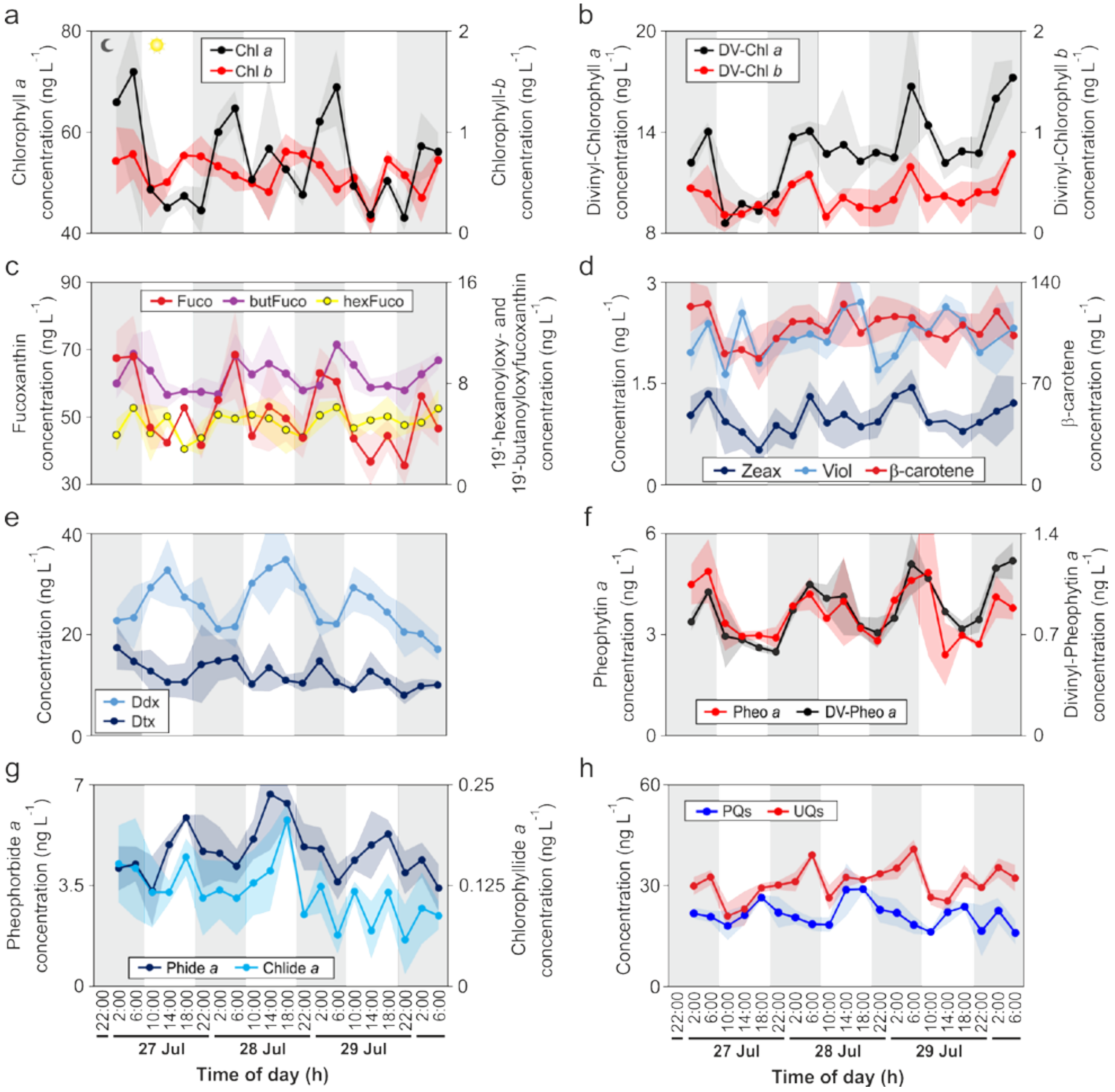


**Figure S1.** Diel variations of concentrations of **a** Chl *a* and *b*, **b** DV-Chl *a* and *b*, **c** Fuco, butFuco, and hexFuco, **d** Zeax, Viol, and β-carotene, **e** Ddx and Dtx, **f** Pheo *a* and DV-Pheo *a*, **g** Phide a and Chlide *a*, **h** PQs and UQs over 4 day/night cycles at 15m water depth in July and August 2015 at Station ALOHA. Values represent the average of environmental triplicates and shaded envelope the standard deviation (n=3). Gray bars indicates night while white indicates day. Chl, chlorophyll; DV-Chl, divinyl chlorophyll; Fuco, fucoxanthin; butFuco, 19'-butanoyloxyfucoxanthin; hexFuco, 19'-hexanoyloxyfucoxanthin; Zeax, zeaxanthin; Viol, violaxanthin; Ddx, diadinoxanthin; Dtx, diatoxanthin; Pheo, pheophytin; DV-Pheo, divinyl pheophytin; Phide, pheophorbide; Chlide, chlorophyllide; PQs, plastoquinones; UQs, ubiquinones.

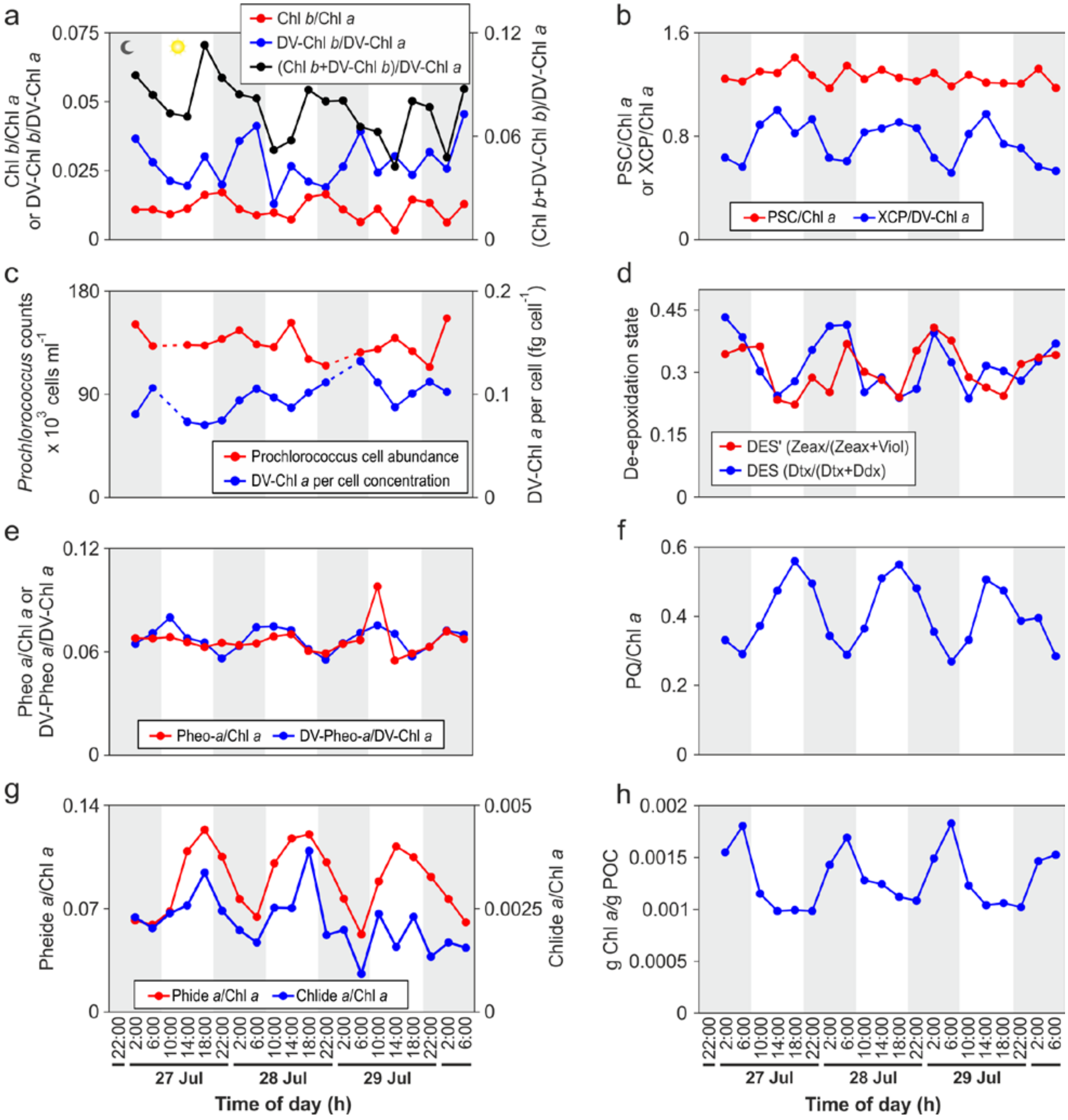


**Figure S2.** Diel variations of **a** Chl *b*/Chl *a*, DV-Chl *b*/DV-Chl *a* and (Chl *b*+DV-Chl *b*)/DV-Chl *a* **b** PSC/Chl *a* and XCP/Chl *a*, **c** *Prochlorococcus* cell abundance and DV-Chl a per cell concentrations, **d** De-epoxidation states of xanthophyll cycle pigments, **e** Pheo *a*/Chl *a* and DV-Pheo *a*/DV-Chl *a*, **f** PQ/Chl *a*, **g** Pheide *a*/Chl *a* and Chlide *a*/Chl *a*, and **h** g Chl *a*/g POC over 4 day/night cycles at 15m water depth in July and August 2015 at Station ALOHA. Chl, chlorophyll; DV-Chl, divinyl chlorophyll; PSC, photosyntheitc pigments; XCP, xanthophyll cycle pigents; Zeax, zeaxanthin; Viol, violaxanthin; Ddx, diadinoxanthin; Dtx, diatoxanthin; DES, de-epoxidation state; Pheo, pheophytin; DV-Pheo, divinyl pheophytin; Phide, pheophorbide; Chlide, chlorophyllide; PQs, plastoquinones.

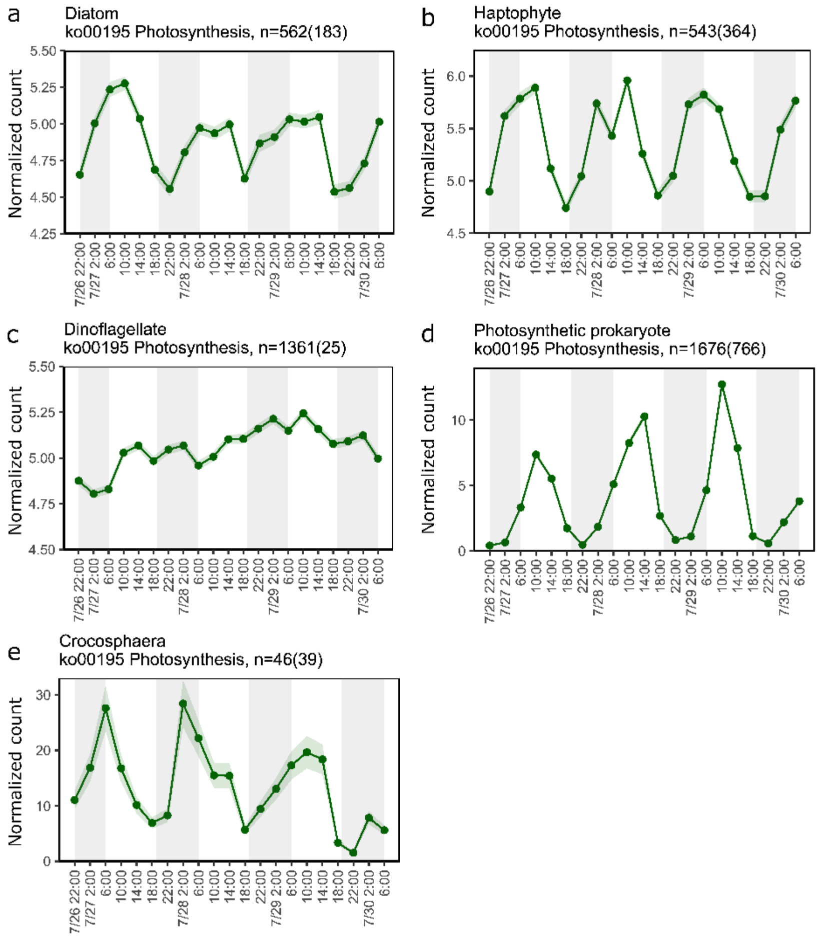


**Figure S3**. Averaged normalized expression patterns for genes within the KEGG Photosynthesis pathway (ko00195) across the sampling period for a) diatoms, b) haptophytes, c) dinoflagellates, d) photosynthetic prokaryotes, and e) *Crocosphaera*. Shaded envelope is the standard error of the mean. Gray bars indicates night while white indicates day. The number of genes represented within each figure is displayed in the heading with the number of significantly periodic (RAIN, FDR ≤0.05) genes in parentheses.

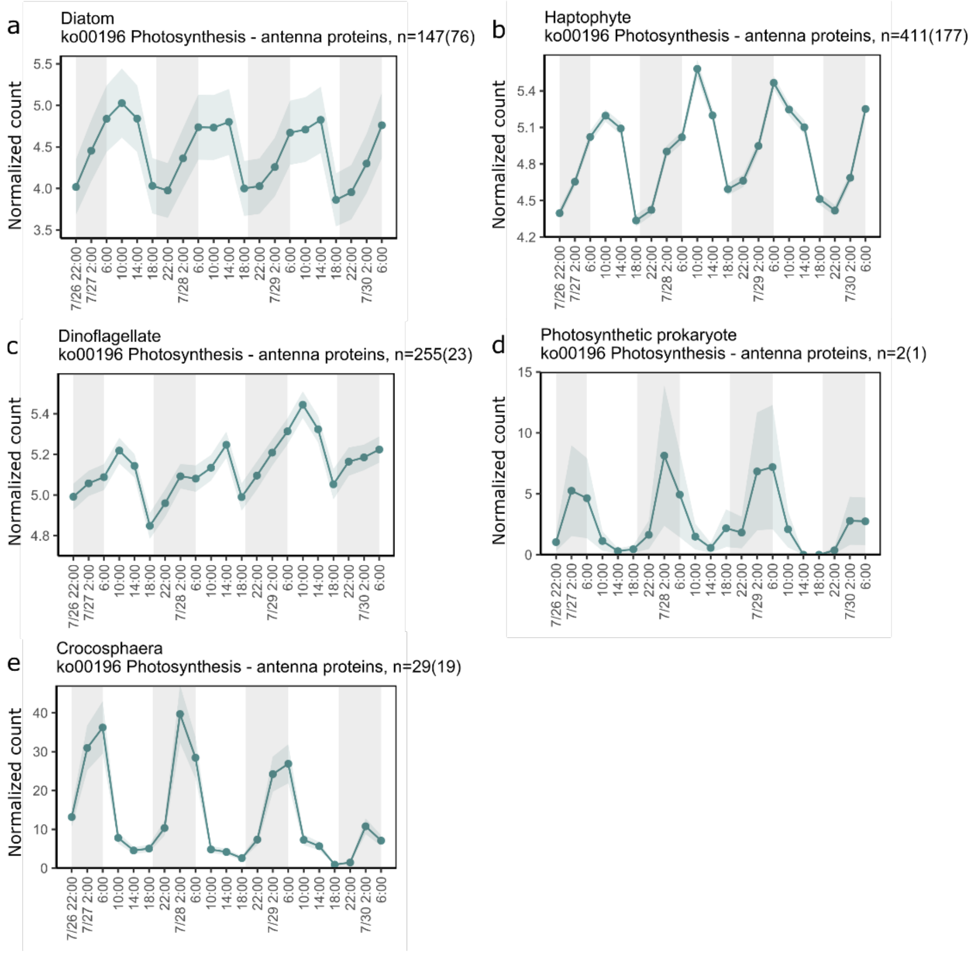


**Figure S4**. Averaged normalized expression patterns for genes within the KEGG Photosynthesis - antenna proteins pathway (ko00196) across the sampling period for **a** diatoms, **b** haptophytes, **c** dinoflagellates, **d** photosynthetic prokaryotes, and **e** *Crocosphaera*. Shaded envelope is the standard error of the mean. Gray bars indicates night while white indicates day. The number of genes represented within each figure is displayed in the heading with the number of significantly periodic(RAIN, FDR ≤0.05) genes in parentheses.

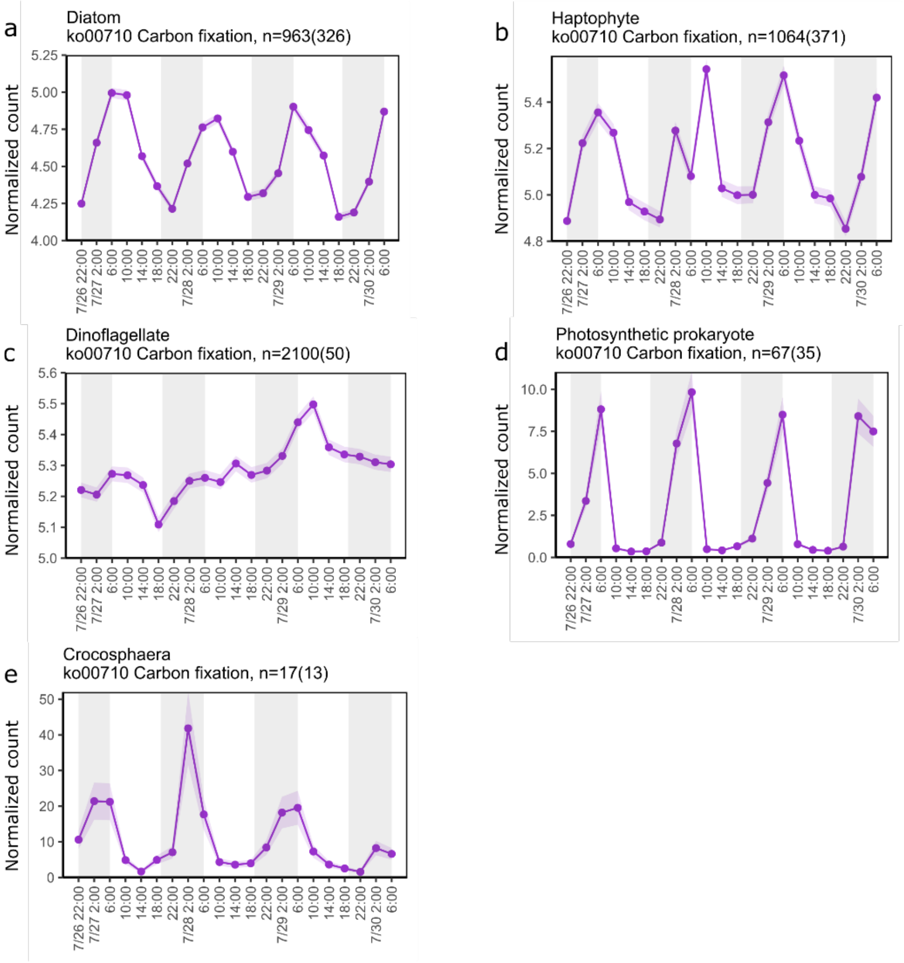


**Figure S5**. Averaged normalized expression patterns for genes within the KEGG Carbon fixation in photosynthetic organisms pathway (ko00710) across the sampling period for **a** diatoms, **b** haptophytes, **c** dinoflagellates, **d** photosynthetic prokaryotes, and **e** *Crocosphaera*. Shaded envelope is the standard error of the mean. Gray bars indicates night while white indicates day. The number of genes represented within each figure is displayed in the heading with the number of significantly periodic (RAIN, FDR ≤0.05) genes in parentheses.

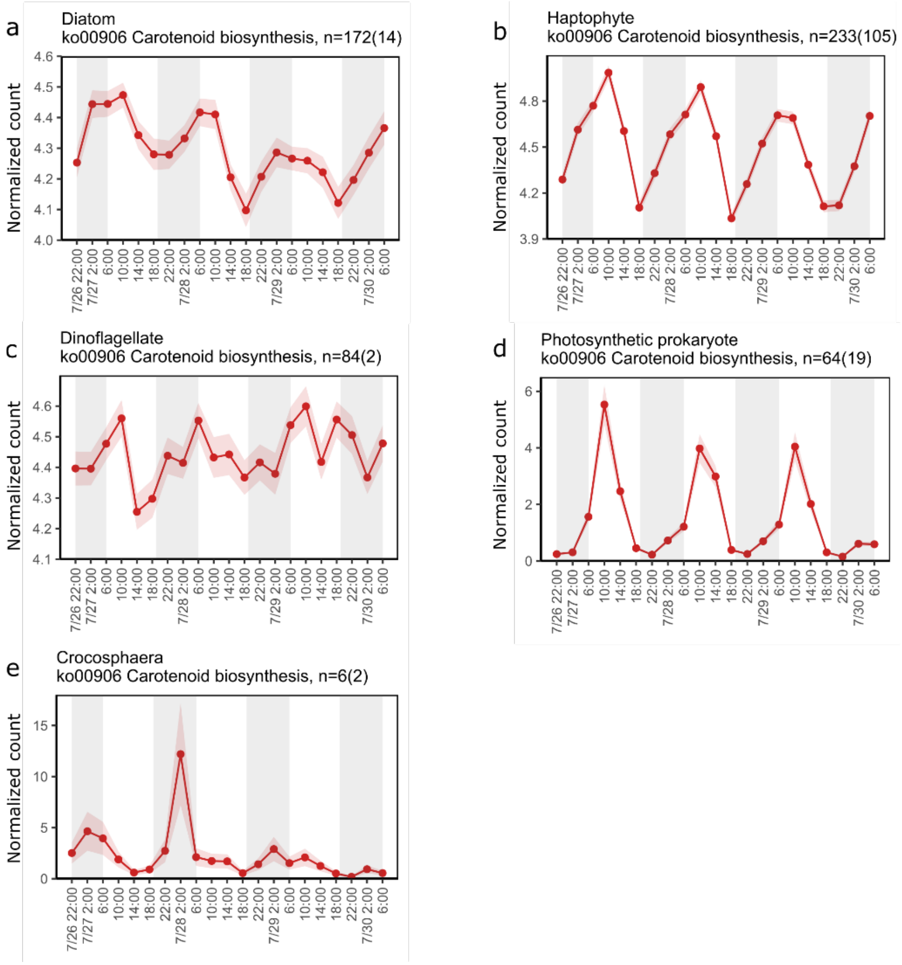


**Figure S6**. Averaged normalized expression patterns for genes within the KEGG Carotenoid biosynthesis pathway (ko00906) across the sampling period for **a** diatoms, **b** haptophytes, **c** dinoflagellates, **d** photosynthetic prokaryotes, and **e** *Crocosphaera*. Shaded envelope is the standard error of the mean. Gray bars indicates night while white indicates day. The number of genes represented within each figure is displayed in the heading with the number of significantly periodic (RAIN, FDR ≤0.05) genes in parentheses.

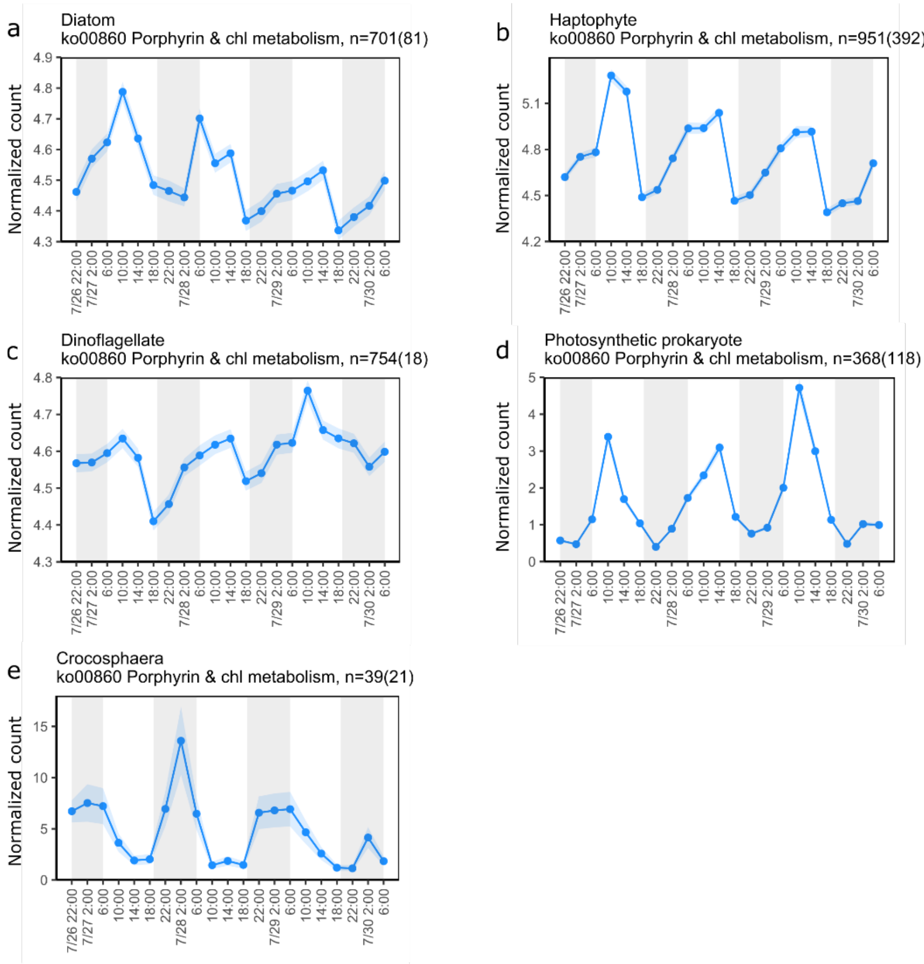


**Figure S7**. Averaged normalized expression patterns for genes within the KEGG Porphyrin and chlorophyll metabolism pathway (ko00860) across the sampling period for **a** diatoms, **b** haptophytes, **c** dinoflagellates, **d** photosynthetic prokaryotes, and **e** *Crocosphaera*. Shaded envelope is the standard error of the mean. Gray bars indicates night while white indicates day. The number of genes represented within each figure is displayed in the heading with the number of significantly periodic (RAIN, FDR ≤0.05) genes in parentheses.

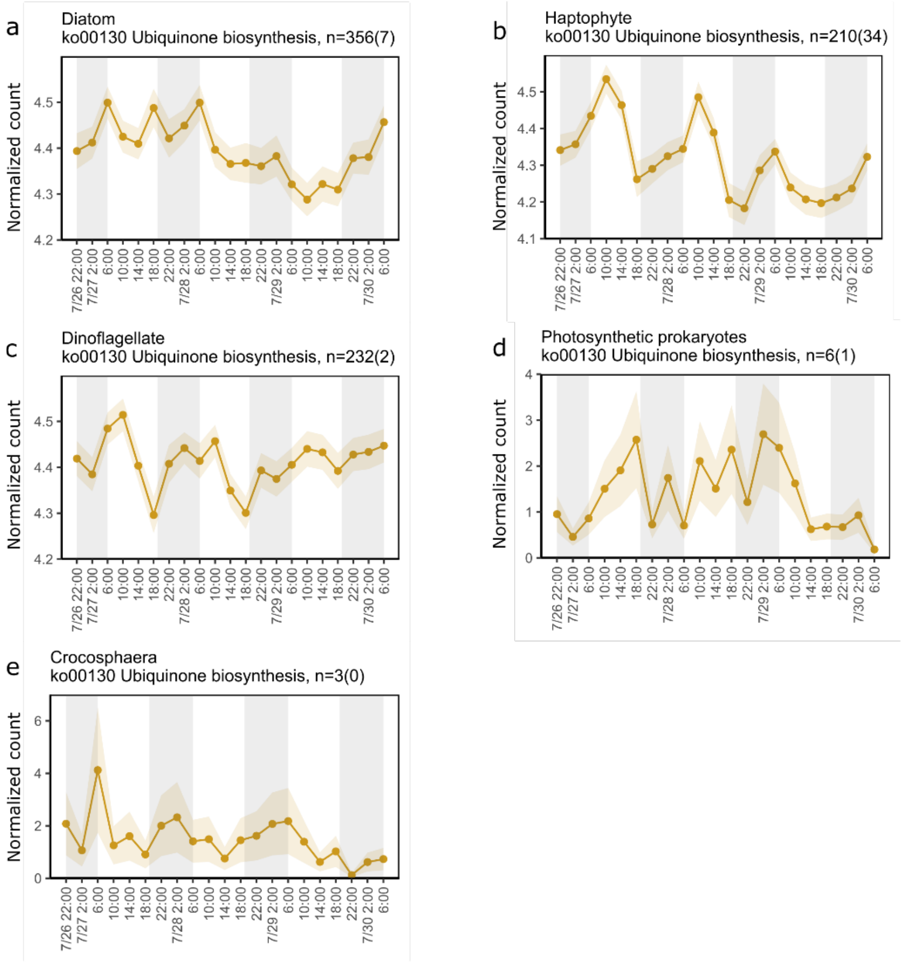


**Figure S8**. Averaged normalized expression patterns for genes within the KEGG Ubiquinone and other terpenoid-quinone biosynthesis pathway (ko00130) which are involved in ubiquinone biosynthesis across the sampling period for **a** diatoms, **b** haptophytes, **c** dinoflagellates, **d** photosynthetic prokaryotes, and **e** *Crocosphaera*. Shaded envelope is the standard error of the mean. Gray bars indicates night while white indicates day. The number of genes represented within each figure is displayed in the heading with the number of significantly periodic (RAIN, FDR ≤0.05) genes in parentheses.

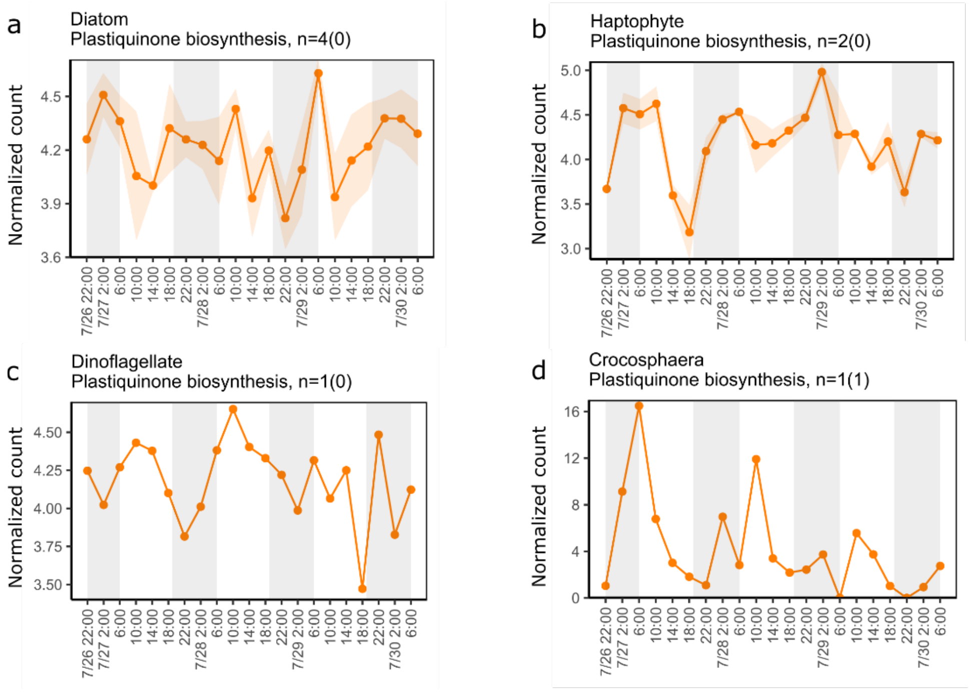


**Figure S9**. Averaged normalized expression patterns for genes within the KEGG Ubiquinone and other terpenoid-quinone biosynthesis pathway (ko00130) which are involved in plastiquinone biosynthesis across the sampling period for **a** diatoms, **b** haptophytes, **c** dinoflagellates, **d** *Crocosphaera*. Shaded envelope is the standard error of the mean. Gray bars indicates night while white indicates day. The number of genes represented within each figure is displayed in the heading with the number of significantly periodic (RAIN, FDR ≤0.05) genes in parentheses.

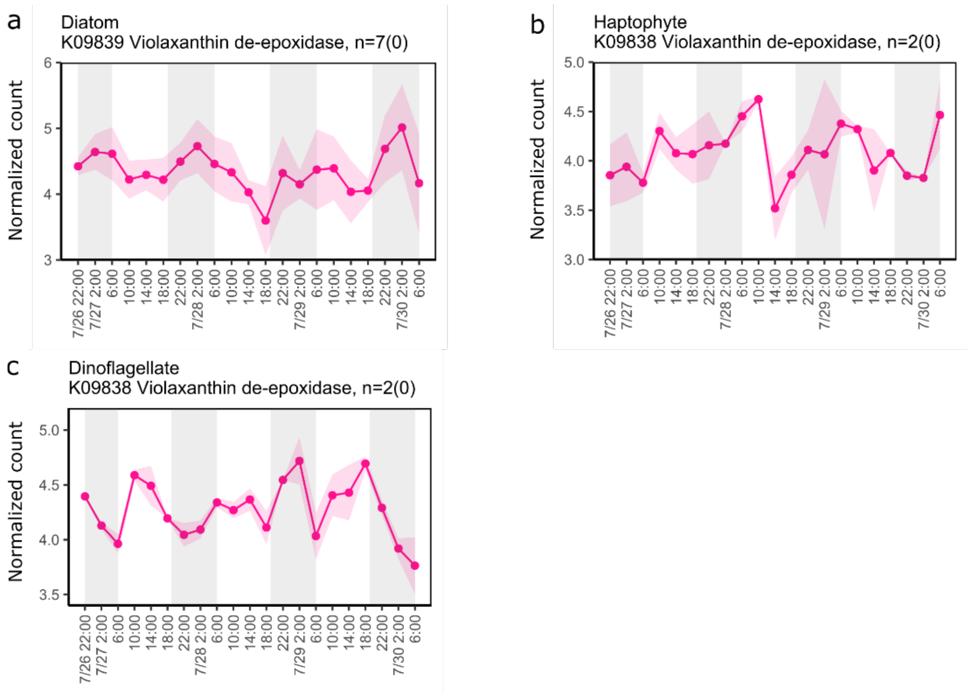


**Figure S10**. Averaged normalized expression patterns for genes involved in xanthophyll de-epoxidation across the sampling period for **a** diatoms, **b** haptophytes, and **c** dinoflagellates. Shaded envelope is the standard error of the mean. Gray bars indicates night while white indicates day. The number of genes represented within each figure is displayed in the heading with the number of significantly periodic (RAIN, FDR ≤0.05) genes in parentheses.

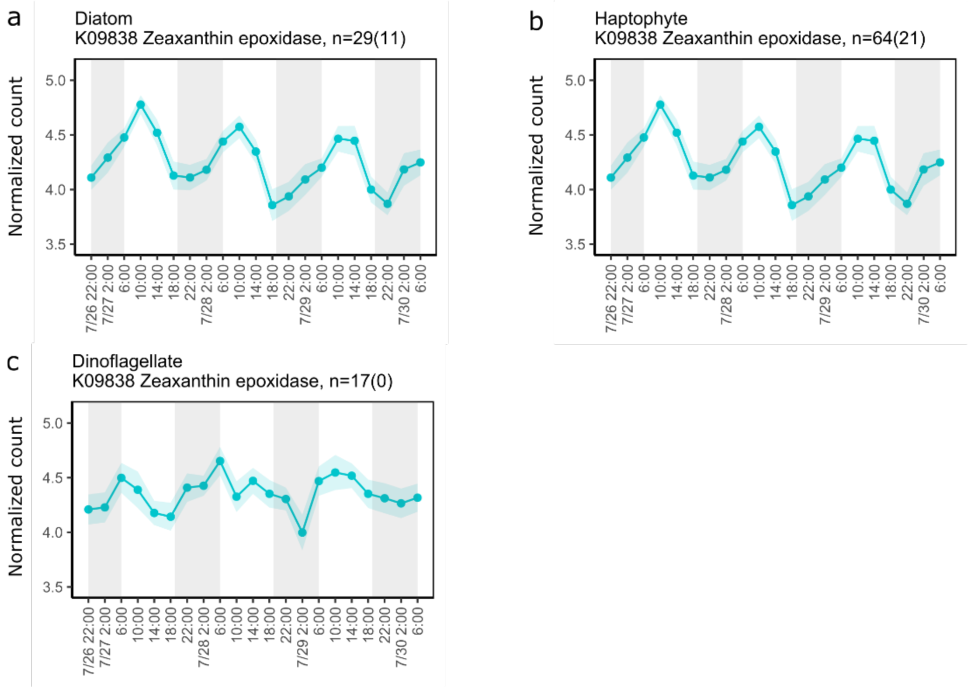


**Figure S11**. Averaged normalized expression patterns for genes involved in xanthophyll epoxidation across the sampling period for **a** diatoms, **b** haptophytes, and **c** dinoflagellates. Shaded envelope is the standard error of the mean. Gray bars indicates night while white indicates day. The number of genes represented within each figure is displayed in the heading with the number of significantly periodic (RAIN, FDR ≤0.05) genes in parentheses.

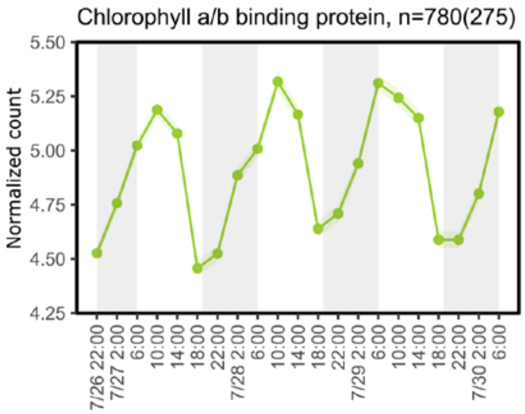


**Figure S12**. Averaged normalized expression patterns of chlorophyll a/b binding protein genes (ko08907-08917 and ko14172) in eukaryotes. Shaded envelope is the standard error of the mean. Gray bars indicates night while white indicates day. The number of genes is displayed in the heading with the number of significantly periodic (RAIN, FDR ≤0.05) genes in parentheses. K08907-8917 and K14172.

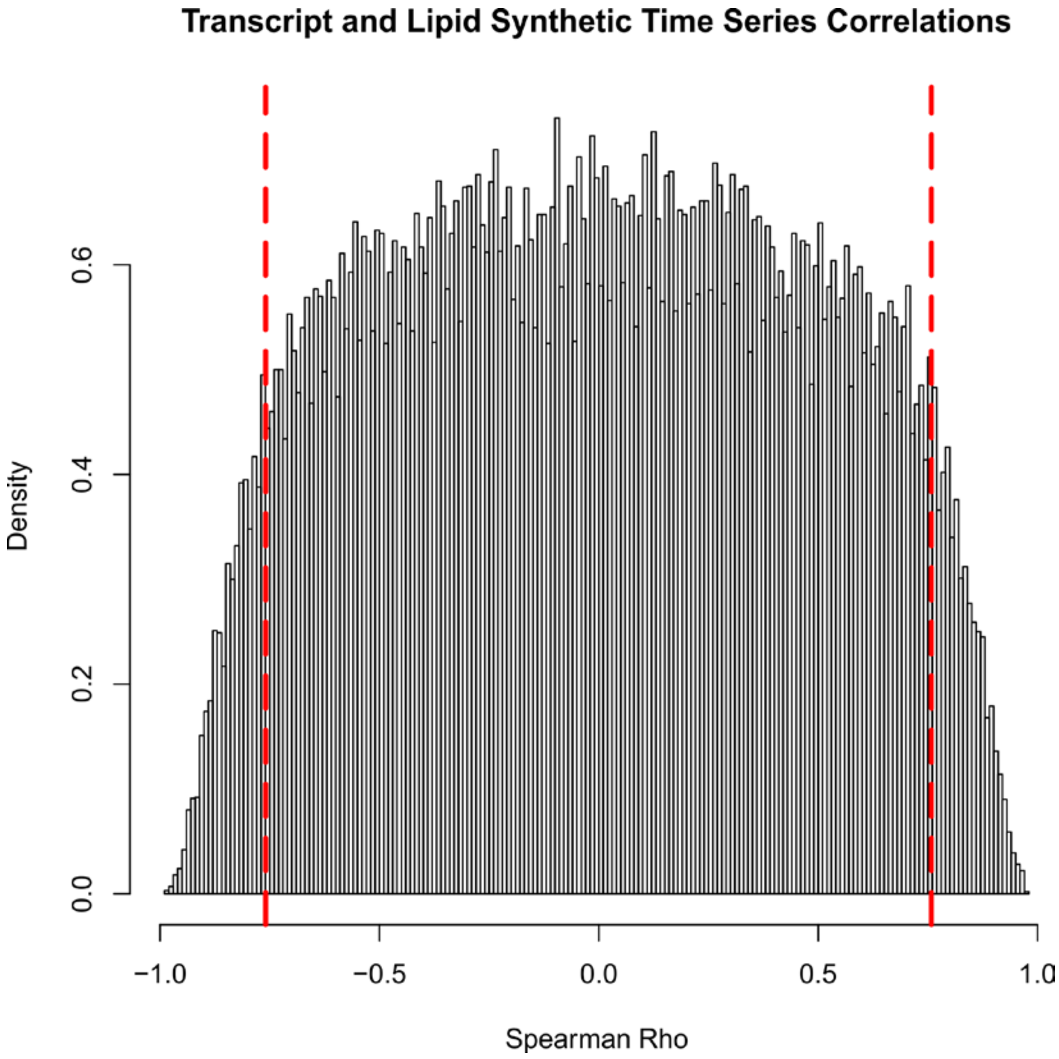


**Figure S13**. Histogram of simulated spearman's rho values for independently generated synthetic time series. 100,000 time series were simulated by bootstrap sampling from the distribution of observed lag 1 differences in transcript and lipid time series and cumulatively summing them. The spearman's rho was then calculated between 100,000 randomly chosen pairs from amongst these synthetic time series to simulate the probability density function of spearman's rho values between iid random walks (shown here) using the distribution of step sizes from the data. Red dashed lines indicate the 5th and 95th percentiles.
